# Cytokine profiles in adults with imported malaria: insights from the PALUREA cohort study

**DOI:** 10.1101/2022.11.23.517616

**Authors:** Charles de Roquetaillade, Cédric Laouenan, Jean-Paul Mira, Carine Roy, Marie Thuong, Élie Azoulay, Didier Gruson, Frédéric Jacobs, Juliette Chommeloux, François Raffi, Laurent Hocqueloux, Patrick Imbert, Vincent Jeantils, Jean-Louis Delassus, Sophie Matheron, Catherine Fitting, Jean-François Timsit, Fabrice Bruneel

## Abstract

The increase in worldwide travel is making imported malaria a growing health concern in nonendemic countries. Most data on the pathophysiology of malaria come from endemic areas. Little is known about cytokine profiles during imported malaria. We report cytokine profiles in adults with *Plasmodium falciparum* malaria included in PALUREA, a prospective cohort study conducted in France between 2006 and 2010. The patients were classified as having uncomplicated malaria (UM) or severe malaria (SM), with this last further categorized as very severe malaria (VSM) or less severe malaria (LSM). At hospital admission, eight blood cytokines were assayed in duplicate using Luminex technology: interleukin (IL)-1α, IL-1β, IL-2, IL-4, IL-10, tumor necrosis factor (TNF)α, interferon (IFN)γ, and macrophage migration inhibitory factor (MIF). These assays were repeated on days 1 and 2 in the SM group. Of the 278 patients, 134 had UM and 144 SM. At hospital admission, over half the patients had undetectable levels of IL-1α, IL-1 β, IL-2, IL-4, IFN γ, and TNFα, while IL-10 and MIF were significantly higher in the SM vs. the UM group. Higher IL-10 was significantly associated with higher parasitemia (R=0.32 [0.16–0.46]; *P*=0.0001). In the SM group, IL-10 elevation persisting from admission to day 2 was significantly associated with subsequent nosocomial infection. Of eight tested cytokines, only MIF and IL-10 were associated with disease severity in adults with imported *P. falciparum* malaria. At admission, many patients had undetectable cytokine levels, suggesting that circulating cytokine assays may not be helpful as part of the routine evaluation of adults with imported malaria.

**Author Summary:** *Plasmodium falciparum* malaria is increasingly common in nonendemic areas. Improved understanding of its pathophysiology might help to decrease mortality. We therefore routinely assayed eight cytokines in 278 adults with imported *P. falciparum* malaria at hospital admission; in the group with severe malaria (n=144), we repeated the assays on the next two days. The cytokine levels were often undetectable, suggesting that cytokine storm might not be a driving mechanism at the time of clinical presentation. IL-10 and macrophage migration inhibitory factor (MIF) were significantly higher in the group with severe vs. uncomplicated disease. Thus, the roles for these two cytokines in severe malaria, may deserve further investigation. A complicating factor is that greater IL-10 elevation may be a response to a heavy parasite burden and/or may promote parasite replication. IL-10 elevation that persisted over the first 2 days after admission was significantly associated with subsequent nosocomial infections in the group with severe malaria suggesting its possible role in acquired immune suppression syndrome.

## INTRODUCTION

*Plasmodium falciparum* malaria has been a global health priority for over 30 years. It caused 627 000 deaths in 2020(1). Imported malaria, defined as contracted in an endemic area but developing in a non-endemic country, is a growing health concern in the northern hemisphere due to the increase in worldwide travel. Among European countries, France has the most cases, about 5000 annually, of which 10%–15% are severe and the great majority are due to *P. falciparum*. In France, severe imported malaria was fatal in 3% to 11% of patients in recent studies(2–4). Given the available medical resources, lower rates should be achievable(2).

Most data on the pathophysiology of falciparum malaria come from children in endemic areas(5). However, we have previously reported that patients with imported malaria differ from those in endemic areas. Most are adults, usually Caucasian, and with no past *P. falciparum* exposure(4). Importantly, despite better baseline health and greater access to healthcare resources, severe forms are more common. These differences may be related to host factors that affect the immune response. We know little about the immune response to imported malaria. The few studies of the cytokine signature during imported malaria were done in small and heterogeneous populations (6,7).

*P. falciparum* infection triggers strong inflammatory and immune responses similar to those seen in bacterial infections(6,8,9) despite the many differences in underlying pathophysiology. Endothelial sequestration of parasitized erythrocytes is unique to malaria and may cause the main manifestations of severe forms. The relative roles for inflammation vs. immune responses in promoting erythrocyte sequestration remain debated.

The primary objective of this study was to describe the cytokine signature in adults with imported falciparum malaria. The secondary objective was to investigate potential associations linking plasma cytokine levels to patient outcomes.

## PATIENTS AND METHODS

### Study design and patients

The present study is a pre-specified ancillary analysis of data from the previously reported prospective multicenter PALUREA cohort study conducted in France between November 2006 and March 2010 and designed to investigate several host- and parasite-related biomarkers, with the objective of improving the evaluation of severity(10). The study protocol was approved for all participating centers on May 2, 2006, by the ethics committee of the Saint-Louis University Hospital, Paris, France (CCPPRB, approval #2006/24). PALUREA complied with French regulations, the Declaration of Helsinki, and Good Clinical Practices. Written informed consent was obtained before inclusion from each patient, or next of kin if the patient had lost competency; according to French law, incompetent patients with no available next of kin were included then asked, as soon as they regained competency, whether they consented to stay in the study.

As described in detail elsewhere(10), the participating centers enrolled consecutive patients with either uncomplicated malaria (UM) or severe malaria (SM). We defined SM as *P. falciparum* malaria requiring admission to the intensive care unit (ICU) and fulfilling the modified 2000 WHO criteria for severe malaria in adults(11,12) at admission or within the first 2 ICU days (see **Supplementary Table S1**). In the SM group, based on French guidelines(13), we predefined two subgroups, namely, very severe malaria (VSM) and less severe malaria (LSM). VSM was defined as any of the following: coma, shock, acidosis, hyperlactatemia >5 mmol/L, or respiratory distress, within the first 72 hours after ICU admission. LSM was defined as SM with none of the criteria for VSM. UM was defined as no requirement for ICU admission and absence of criteria for SM; Patients were managed according to international guidelines.

### Cytokine assays

We assayed a panel of proinflammatory and antiinflammatory cytokines in blood samples collected in 5-mL heparin tubes, at hospital admission (day 0) in all patients and on days 1 and 2 in patients with SM. The tubes were transferred to the Cochin University Hospital (Paris, France) within 2 hours, in cooled bags, then centrifuged for 10 minutes at 3500 rpm at 4 °C to allow plasma collection. The following eight cytokines were assayed in duplicate using Luminex technology (Thermo Fisher®, Waltham, MA): interleukin (IL)-1α (IL-1α), IL-1β, IL-2, IL-4, IL-10, tumor necrosis factor (TNF)-α, interferon (IFN) -γ, and macrophage migration inhibitory factor (MIF). In patients with SM, the same cytokines were assayed using the same method on days 1 and 2.

### Statistics

In results, continuous data are described as median and interquartile range [IQR], and qualitative variables as number and percentage. Cytokine values were also described with boxplots according to the different patient groups.

Comparisons between two groups (UM vs SM or LSM vs VSM) were performed with Student t-test or Mann-Whitney test as appropriate for continuous variables, and with chi-square test or Fisher exact test as appropriate for categorical variables. To compare plasmatic cytokines level between the three groups UM, LSM and VSM, the non-parametric Kruskal-Wallis test was used.

The association between plasmatic cytokines level and some of biological data was measured using spearman correlation coefficients.

All statistical analyses were performed using SAS version 9.4 software (SAS Institute Inc., Cary, NC).

## RESULTS

### Clinical characteristics (Table 1)

#### Baseline characteristics

**Supplementary Figure 1** is the patient flowchart. Of the 278 patients with cytokine data, 134 had UM and 144 SM, including 68 with VSM and 76 with LSM. **Table 1** reports their baseline characteristics. Briefly, the SM group was characterized by older age, a higher proportion of patients with comorbidities, a longer time from symptom onset to hospital admission, higher parasitemia on day 0, and a higher *pf*HRP2 level. Coinfection at admission was present in 10 SM patients vs. a single UM patient. The rate of adherence to antimalarial chemoprophylaxis was not different between the two groups. Nosocomial infections were significantly more common in the SM group. All 8 patients who died had SM (8/144, 5.5%).

#### Patients with very severe malaria vs. less severe malaria

**Table 1** compares the VSM and LSM groups. VSM patients were older and less likely to be of African origin. Time from symptom onset to hospital admission was not different between the two sub-groups. The higher *pf*HRP2 value in the VSM sub-group despite no significant parasitemia difference with the LSM sub-group on day 0 suggested a greater sequestered parasite biomass(14).

#### Circulating cytokine levels at hospital admission (day 0) (Figure 1 and Table 2)

**Figure 1.**
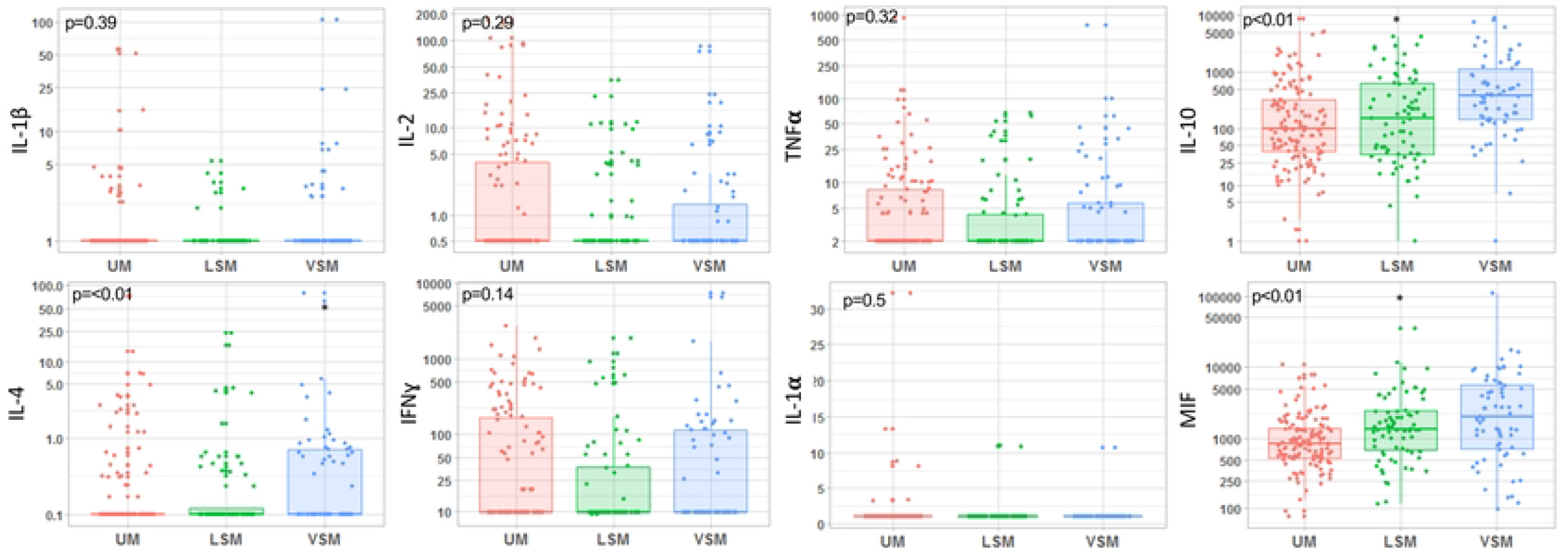
Boxplot of circulating cytokine levels at admission (D0) in patients with uncomplicated malaria (UM), less severe malaria (LSM), and very severe malaria (VSM). Kruskal-Wallis test for intergroup distribution, with *P* values <0.05 considered significant

**Figure 1** reports the day 0 cytokine levels in the UM group and LSM and VSM sub-groups. Levels were often undetectable for six cytokines (IL-1 α, IL-1β, IL-2, IL-4, IFNγ, and TNFα) (**Table 2**). IL-4 was more often detectable in the SM group than in the UM group(*P*=0.04); for the other cytokines, the proportions of patients with detectable levels were not different across groups and sub-groups (**Table 2**). IL-4, IL-10, and MIF were significantly higher in patients with greater disease severity (**Figure 1**). Only IL-10 was significantly associated with parasitemia (R=0.32 [0.16–0.46]; *P*=0.0001). IL-4, IFNγ, TNFα, IL-10, and MIF were significantly but only weakly associated with the *pf*HRP2 level at admission (**Supplementary Table S2**). None of the cytokine levels were associated with albuminemia or hemoglobin level at hospital admission. The cytokine profile at admission did not differ between the patients who died (n=8) and those who survived (**Supplementary Table S3**).

#### Changes in circulating IL-10 and MIF levels during the first 3 ICU days (Figure 2)

In the 144 patients with SM, the eight cytokines were assayed on the first ICU Day then on the next 2 days. Median IL-10 levels were significantly higher in the VSM vs. the LSM sub-group on days 1 and 2 (day 1: 209.36 [58.44–693.21] vs. 72.49 [35.01–169.52], *P*<0.01; day 2: 195.20 [52.34–532.60] vs. 37.78 [15.20–151.35] on day 2, *P*<0.0; **Figure 2**). The median MIF values were significantly higher in the VSM sub-group vs. the LSM sub-group on day 2 but not on day 1 (day 1: 1676.0 ng/mL [898.16–3321.60] vs. 1206.90 ng/mL [507.98–2960.50], *P*=0.12; day 2: 1462.20 ng/mL [825.94–4898.10] vs. 1083.20 ng/mL [631.91–2687.50], *P*=0.02). Interestingly, high IL-10 levels persisting over the 3 days were more common in patients with vs. without nosocomial infections (**Supplementary Table S4**).

**Figure 2.**
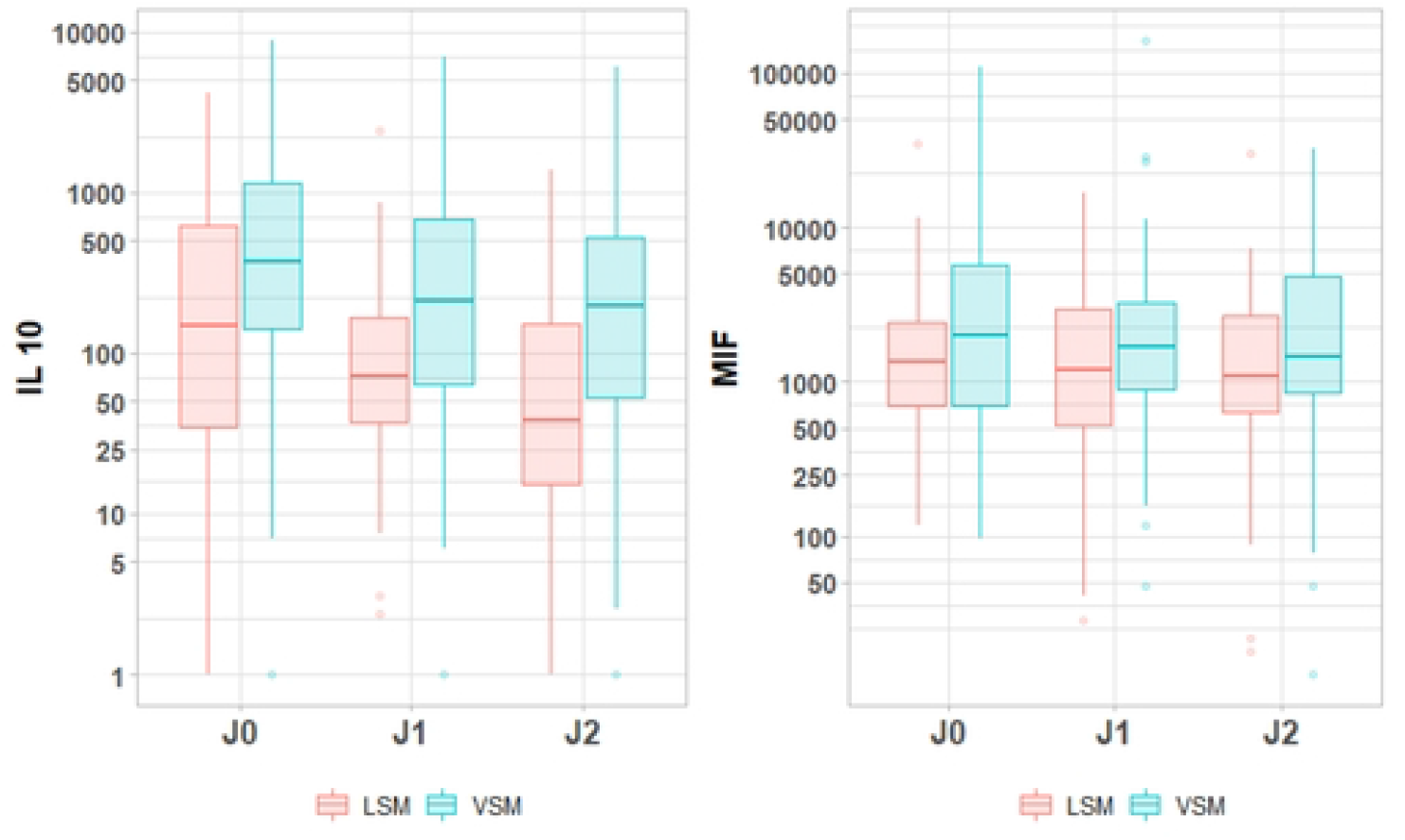
Boxplot of circulating cytokine levels in the sub-groups with less severe malaria (LSM) and very severe malaria (VSM) during the first 3 days following ICU admission. Sub-group comparison using the t-test between groups, with *P* values <0.05 considered significant

## DISCUSSION

In this preplanned analysis of data from the prospective PALUREA observational cohort study [8], we evaluated circulating levels of eight cytokines in 278 adults admitted for imported malaria, including 134 with UM and 144 with SM. At hospital admission, IL-1α, IL-1β, IL-2, IL-4, IFNγ, and TNFα were undetectable in over half the patients, whereas IL-10 and MIF levels were detectable and significantly higher in the SM group. Higher IL-10 levels were significantly associated with higher parasitemia.

Cytokines play a pivotal role in the host response to *P. falciparum* infection. We found no meaningful associations with disease severity and frequently undetectable levels for six of the eight tested cytokines (IL-1α, IL-1β, IL2, IL4, TNFα, and IFNγ). It is worth noting that the cytokines were assayed routinely at hospital admission and that their levels may have been influenced by time since symptom onset, whose median value was 4 [2-6] days. For some cytokines such as IFNγ, peak levels may be achieved very early, at the liver stage of malaria, i.e., before symptom onset(15). In an in vitro study, TNFα and IL-1β peaked within 2 h after stimulation with hemozoin (16). Thus, the assays in our study may have been done after the peak for some cytokines. A technical issue as the reason for the undetectable cytokine levels is unlikely, given the duplicate assays at a central, highly experienced laboratory. The most reasonable interpretation is that levels of the tested cytokines were low in most patients at the time of inclusion.

A good balance between the proinflammatory and anti-inflammatory host responses is crucial to parasite eradication without further organ damage due to runaway inflammation. IL-10 is a fundamental anti-inflammatory cytokine that plays an important regulatory function in malarial models(17,18). In our study, IL-10 was associated with severity as found in several studies(7,19–22). However, the precise role of IL-10 in malaria is unclear. On one hand, one study found a protective effect of IL-10 by inhibition of endothelial ICAM-1 expression, resulting in a decrease in parasite sequestration(23). On another hand, due to its potent anti-inflammatory effects, IL-10 may promote parasite growth(24). The direction of the association between IL-10 and parasite burden cannot be determined from our data: IL-10 elevation may have been triggered by a heavy parasite burden and/or parasite replication may have been promoted by high IL-10 levels. Finally, the strong immunosuppressive effect of IL-10 on circulating monocytes(25,26) may lead to increased susceptibility to bacterial infections(27). In our group with SM, patients with high IL-10 levels during the first 3 ICU days had a significantly greater number of nosocomial infections.

MIF, one of the first cytokines identified, is constitutively expressed by a broad spectrum of cells and tissues. MIF contributes to regulate immune responses and is an important mediator of inflammatory diseases in mammals(28). Recently, the discovery of a *P. falciparum*-encoded MIF ortholog (PfMIF) drew attention to the role for MIF in the pathophysiology of malaria(29,30). MIF exerts pro-inflammatory properties that may increase the cytoadherence of parasitized erythrocytes, resulting in greater disease severity. MIF also inhibits erythroid, multipotential, and granulocyte–macrophage progenitor-derived colony formation and may therefore be involved in the pathophysiology of malarial anemia(31). We found that higher MIF levels were associated with greater disease severity, in keeping with findings in Zambian children(21) and Indian infants and adults (32). In contrast to the work from Zambia, our study showed no association between MIF levels and anemia. However, the absence of PfMIF assays in our study precludes conclusions about the role for MIF in malarial anemia. To our knowledge, no studies have investigated the levels of both MIF isoforms (human MIF and PfMIF) and their relationship with malaria severity. Therapeutic interventions designed to neutralize PfMIF have shown promise for protecting against malaria(33,34).

The main benefits expected from assessing immune responses in critically ill patients with malaria are the characterization of specific immune dysfunctions and the identification of patients at high risk for nosocomial infections. Higher IL-10 levels may be associated with worse immune dysfunction and greater susceptibility to nosocomial infections. In preclinical studies, high IL-10 levels were associated with decreases in the release of other cytokines and in class II major histocompatibility complex expression on monocytes(35). In our population, persistently high IL-10 levels were associated with subsequent nosocomial infections, as previously reported in sepsis but not in malaria(26). Detailed monitoring of both cytokines and membrane receptors associated with immune suppression such as mHLA-DR is a promising approach in sepsis and deserves further investigations in malaria(27).

The strengths of our study include the multicenter recruitment with a large sample size. Nonetheless, the small number of patients who died may have limited our ability to detect statistically significant differences in cytokine profiles between the nonsurvivors and survivors. That the timing of the cytokine assays after symptom onset, i.e., at the blood stage of infection, may have limited our ability to detect cytokine elevation is discussed above. Also mentioned above is the absence in our study of PfMIF assays. Although all our patients were recruited in France, the management of imported malaria in adults followed international recommendations, supporting the applicability of our findings to other countries with similar healthcare resources.

## CONCLUSION

Among the eight cytokines tested in this large cohort of adults with imported falciparum malaria, only MIF and IL-10 were higher in patients with greater disease severity. Our findings suggest that cytokine assays at and early after admission may have a limited role for the diagnosis and assessment of severity. Whether monitoring the potent anti-inflammatory cytokine IL-10 helps to identify patients at high risk for developing nosocomial infection deserves further investigation.

## ACKNOWLEDGMENTS

This work was supported by a research grant from the French Ministry of Health (*Programme Hospitalier de Recherche Clinique*, PHRC, #AOR05007). The sponsor was the public organization *Département de la Recherche Clinique et du Développement, Assistance Publique - Hôpitaux de Paris*. The trial is registered on ClinicalTrials.gov (#NCT00372684). We thank all the nurses, physicians, and biologists of the PALUREA Study Group for excellent patient care. We are grateful to Prof. Jean-Marc Cavaillon for scientific advice based on his considerable expertise. We are indebted to the *Centre Hospitalier de Versailles* for editorial assistance.

## DATA AVAILABILITY STATEMENT

Data will be made available upon motivated request formulated to the corresponding author

